# AntiCPs-CompML: A Comprehensive Fast Track ML method to predict Anti-Corona Peptides

**DOI:** 10.1101/2024.06.27.601090

**Authors:** Prem Singh Bist, Sadik Bhattarai, Hilal Tayara, Kil To Chong

## Abstract

This work introduces AntiCPs-CompML, a novel Machine learning framework for the rapid identification of anti-coronavirus peptides (ACPs). ACPs, acting as viral shields, offer immense potential for COVID-19 therapeutics. However, traditional laboratory methods for ACP discovery are slow and expensive. AntiCPs-CompML addresses this challenge by utilizing three primary features for peptide sequence analysis: Amino Acid Composition (AAC), Pseudo Amino Acid Composition (PAAC), and Composition-Transition-Distribution (CTD). The framework leverages 26 different machine learning algorithms to effectively predict potential anti-coronavirus peptides. This capability allows for the analysis of vast datasets and the identification of peptides with hallmarks of effective ACPs. AntiCPs-CompML boasts unprecedented speed and cost-effectiveness, significantly accelerating the discovery process while enhancing research efficiency by filtering out less promising options. This method holds promise for developing therapeutic drugs for COVID-19 and potentially other viruses. Our model demonstrates strong performance with an F1 Score of 92.12% and a Roc AUC of 76% in the independent test dataset. Despite these promising results, we are continuously working to refine the model and explore its generalizability to unseen datasets. Future enhancements will include feature-based and oversampling augmentation strategies addressing the limitation of anti-covid peptide data for comprehensive study, along with concrete feature selection algorithms, to further refine the model’s predictive power. AntiCPs-CompML ushers in a new era of expedited anti-covid peptides discovery, accelerating the development of novel antiviral therapies.

## 1 Introduction

Coronaviruses, like SARS-CoV and COVID-19, pose a constant threat due to their ability to mutate, particularly in the spike protein. This ongoing mutation has spurred a new wave of global research on coronaviruses [1]–[3]. The fight against viral diseases hinges on developing new treatments. Researchers are pioneering methods to predict mutations in viruses like SARS-CoV-2, specifically focusing on spike proteins. This advanced prediction, aided by tools like the Sars-escape Network, allows for the design of more effective vaccines that can anticipate and counteract viral adaptations, preventing them from undermining existing therapies and vaccines [4]–[6].

In addition, current research against coronaviruses like COVID-19 explores various avenues, including drugs and immune therapies. Among these approaches, antiviral peptides (AVPs) or peptide-based therapy hold particular promise. Their success in past outbreaks, like SARS in 2002-2003, suggests they could be a valuable tool in combating COVID-19 [7]. Peptides are emerging as powerful tools for fighting diseases, particularly those with challenging targets. Their advantages include the ability to disrupt protein interactions and their faster development timeline compared to traditional drugs. Additionally, researchers have successfully designed peptides that effectively inhibit coronaviruses. For instance, Lu et al. [8] created peptides (HR1P and HR2P) that block viral fusion with host cells, while Zhao et al. [9] identified a peptide fragment (P9) that combats both SARS-CoV and MERS-CoV. These examples demonstrate the promising potential of peptides as novel antiviral therapies.

While traditional lab methods exist for finding anticoronavirus peptides (ACVPs), they’re slow and expensive. Researchers are addressing this by developing computational methods. These methods use machine learning (ML) to analyze various features of peptides (like Meta-iAVP) [10] or even deep learning combined with chemical properties (like ENNAVIA) [11]. This allows for faster ACP identification, but current methods still need improvement for real-world use.

This research explores antiviral peptides (AVPs) against coronavirus called as Anti-Corona Virus Peptides (ACVPs) as a promising avenue. Past studies have shown AVPs’ potential against coronaviruses. However, identifying AVPs with specific activity against coronaviruses (ACVPs) is challenging and time-consuming using traditional methods. This work introduced AntiCPs-CompMl, a novel comprehensive method using machine learning to predict ACVPs efficiently. AntiCPs-CompML leverages a powerful and comprehensive technique with the application of LazyPredict [12] to assess the strengths of multiple models for accurate prediction with a maximum accuracy of 91.8%, Area Under Curve-Receiver Operating Characteristics (AUC-ROC) of 0.767 and F1 score of 0.921 for imbalance independent test dataset.

By making ACP discovery faster and more efficient, AntiCPs-CompML paves the way for the development of new coronavirus treatments. This advancement holds immense potential to improve human health and combat future viral threats.

## II. Methods

### A. AntiCPs-CompML Dataset

The AntiCPs-CompML dataset, containing 2,141 entries were retrieved from FEOpti-ACVP [13], serves as a valuable resource for developing and evaluating anti-corona peptide prediction models. It encompasses peptides with varying lengths, ranging from a minimum of 6 amino acids to a maximum of 99. Among these entries, 157 are ACVPs (anticorona virus peptide), while the remainder are non-ACVPs. The dataset is strategically divided, with approximately 80% (1,712 entries) allocated to the training set, ensuring the model encounters a diverse range of examples for robust development. The remaining 20% (429 entries) constitutes the test set, designed to evaluate the model’s generalizability on unseen data. This test set includes 397 non-ACVPs and 32 ACVPs.

### B. Architecture of AntiCPs-CompML

*1) Overview:* The AntiCPs-CompML process encompasses a meticulous workflow: constructing well-defined training and test sets, extracting composition-based informative features from the data, training various machine learning models, conducting independent testing on unseen data, and culminating in a comprehensive performance analysis. The architectural details are illustrated in Figure 1.

**Fig. 1.**
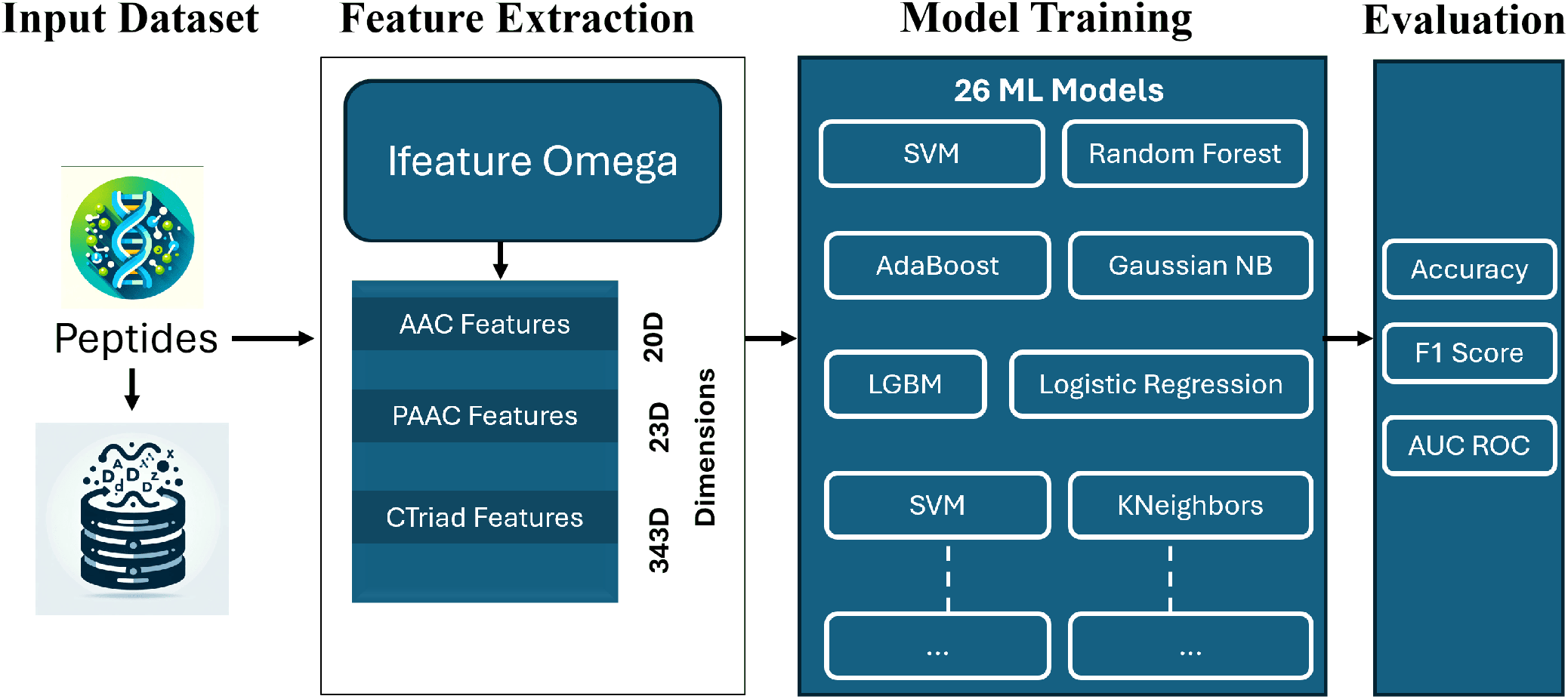
AntiCPs-CompML Workflow:. Feature Extraction and Machine Learning Model Training. The figure illustrates the workflow of the AntiCPs-CompML pipeline for predicting anti-covid peptides (ACPs) from peptide sequences. Informative features are extracted from the peptide sequences using a tool called iFeatureOmega. Twenty-six different machine learning models are trained on the extracted features. The performance of these models is then evaluated using three metrics: Accuracy, ROC AUC score, and F1-Score taking account of imbalance case in the independent test dataset.

### C. Key Feature Extraction

To capture the intricacies of anti-covid peptides (ACVPs), we employed iFeatureOmega [14] for feature extraction from the peptide sequences. This tool facilitated the extraction of informative features categorized into three key groups: Amino Acid Composition (AAC), Composition-Triad (CTriad), and Pseudo Amino Acid Composition (PAAC). These categories were specifically chosen because, compared to others, they have demonstrably proven effective in learning peptide features. Notably, we went beyond individual feature sets by fusing them, creating a more complex feature representation. The feature extraction process yielded informative representations with varying dimensionality: CTriad captured peptide information in 343 dimensions, PAAC in 23 dimensions, and AAC in 20 dimensions. This enriched representation empowers the AntiCPs-CompML model to grasp the nuanced complexities within the peptide sequences.

### D. Harnessing Machine Learning Power

To identify the most effective approach for anti-viral peptide prediction, we leveraged a comprehensive model framework. This framework evaluated the performance of 26 machine learning models, including popular choices like Random Forest, AdaBoost Classifier, Support Vector Machines, K-Nearest Neighbors, and Logistic Regression. Following this comprehensive evaluation, we narrowed our focus to the top ten performing models. These models were strategically chosen based on their Area Under the Curve (AUC) scores, a metric that effectively gauges their ability to distinguish between anti-corona peptides and non-anti-corona peptides.

### E. Performance Measurement

To comprehensively assess the model’s performance in predicting anti-covid peptides, we employed three key evaluation metrics: Accuracy, ROC AUC, and F1-Score. Each metric offers valuable insights: Accuracy measures the overall number of correct predictions, ROC AUC gauges the model’s ability to discriminate between ACPs and non-ACPs, and F1-Score balances precision and recall.

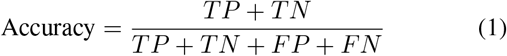

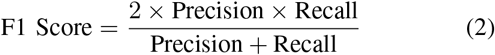

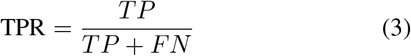

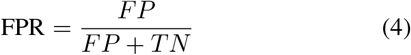

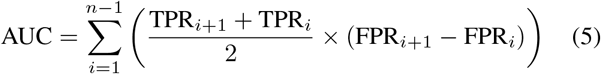

Where TPR_*i*_ and FPR_*i*_ represent the True Positive Rate and False Positive Rate at threshold *i* respectively. AUC is calculated by summing the areas of all trapezoids formed under the ROC curve using *n* different thresholds.

## III. Results

### A. Evaluation of Machine Learning Models for Anti-Covid Peptide Detection

We evaluated the performance of 26 machine learning models for anti-corona peptide (ACVP) detection. The top ten performing models are visualized in Figure 2, Table 1. Each model’s effectiveness was assessed using three key metrics: Accuracy (overall prediction correctness), ROC AUC (ability to discriminate between ACVPs and non-ACVPs), and F1-Score (balance between precision and recall).

**TABLE I.** Anti-covid Peptide (ACP) Detection: Performance Metrics OF 26 Machine Learning Models.

| Model | Accuracy | ROC AUC | F1 Score |
| --- | --- | --- | --- |
| LogisticRegression | 0.918 | 0.767 | 0.921 |
| KNeighborsClassifier | 0.938 | 0.731 | 0.933 |
| PassiveAggressiveClassifier | 0.876 | 0.728 | 0.889 |
| LinearDiscriminantAnalysis | 0.899 | 0.709 | 0.903 |
| BernoulliNB | 0.868 | 0.708 | 0.882 |
| LinearSVC | 0.862 | 0.705 | 0.878 |
| Perceptron | 0.882 | 0.700 | 0.891 |
| RidgeClassifierCV | 0.910 | 0.700 | 0.910 |
| RidgeClassifier | 0.910 | 0.700 | 0.910 |
| SGDClassifier | 0.904 | 0.697 | 0.906 |
| NearestCentroid | 0.859 | 0.688 | 0.875 |
| SVC | 0.932 | 0.680 | 0.924 |
| ExtraTreesClassifier | 0.921 | 0.643 | 0.912 |
| AdaBoostClassifier | 0.899 | 0.631 | 0.895 |
| LGBMClassifier | 0.918 | 0.626 | 0.907 |
| RandomForestClassifier | 0.918 | 0.626 | 0.907 |
| XGBClassifier | 0.915 | 0.624 | 0.905 |
| BaggingClassifier | 0.915 | 0.624 | 0.905 |
| DecisionTreeClassifier | 0.854 | 0.622 | 0.866 |
| GaussianNB | 0.377 | 0.614 | 0.469 |
| ExtraTreeClassifier | 0.876 | 0.603 | 0.878 |
| LabelPropagation | 0.910 | 0.574 | 0.894 |
| LabelSpreading | 0.910 | 0.574 | 0.894 |
| CalibratedClassifierCV | 0.918 | 0.516 | 0.884 |
| DummyClassifier | 0.918 | 0.500 | 0.879 |
| QuadraticDiscriminantAnalysis | 0.907 | 0.494 | 0.874 |

**Fig. 2.**
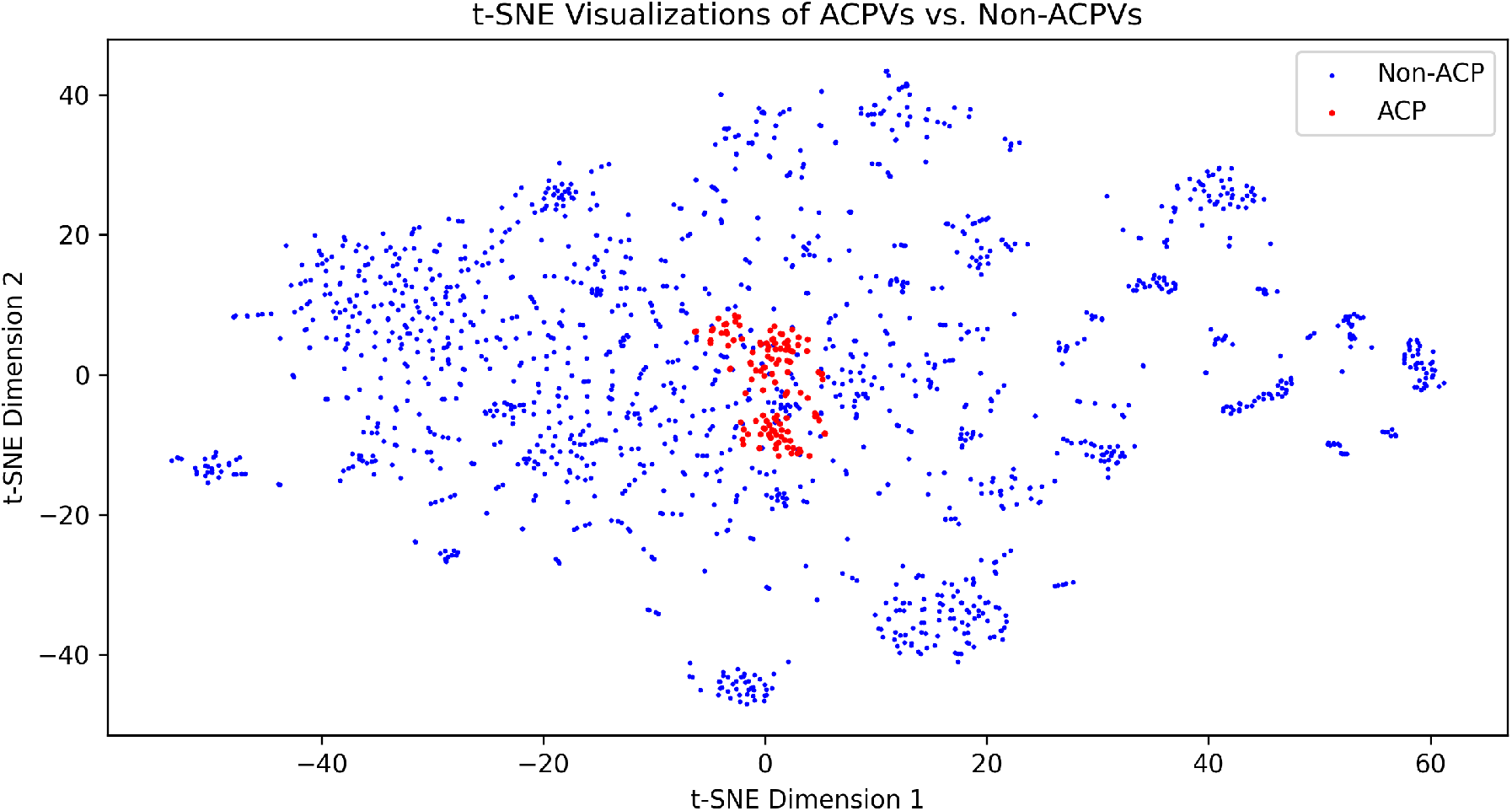
Visualization of ACPs Vs Non-ACPs:. t-SNE visualization of AntiCovid Peptides (ACPs/ACVPs) compared to Non-AntiCovid Peptides (Non-ACPs/Non-ACVPs)

**Fig. 3.**
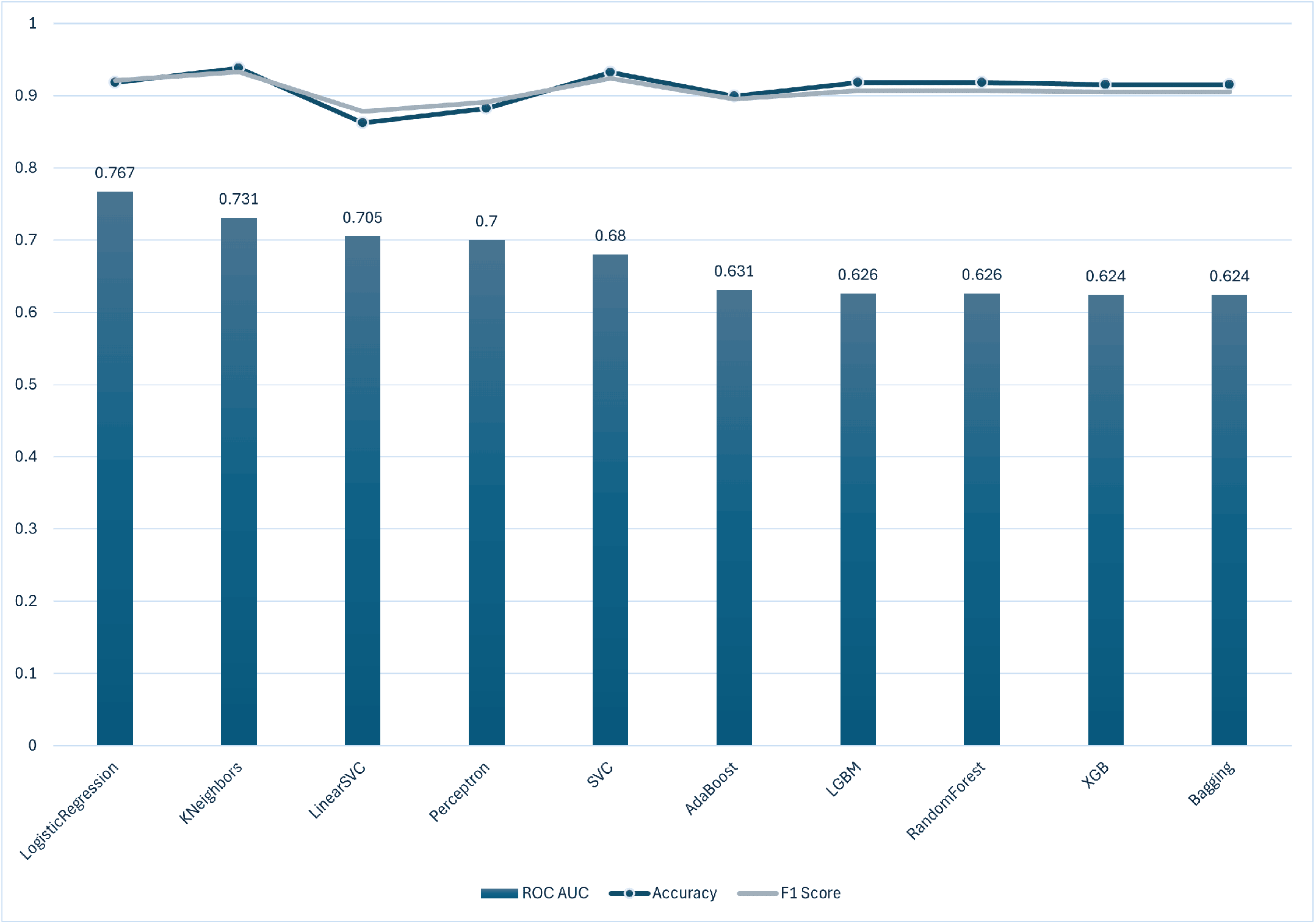
Performance Comparison of Top Ten Machine Learning Models for Anti-Covid Peptide Detection..

Our analysis revealed a fascinating interplay between performance metrics. K-Nearest Neighbors (KNN) emerged as the leader, achieving the highest Accuracy and F1-Score, indicating its prowess in making accurate predictions and identifying true positives. However, Logistic Regression, despite a slight dip in Accuracy and F1-Score, excelled at distinguishing between ACVPs and non-ACVPs, as evidenced by its leading ROC AUC score. This suggests Logistic Regression prioritizes balancing true positive and false positive rates. Linear SVC and Perceptron, while effective, showed a slight decline in all three metrics compared to the top performers. Ensemble methods like AdaBoost, LGBM, Random Forest, XGBoost, and Bagging presented a compelling case with strong Accuracy and F1-Scores, but their lower ROC AUC values suggest they might require further optimization or may be less suitable for datasets with significant class imbalance.

These findings highlight the importance of considering various metrics when selecting a model for ACVP detection. While KNN offers high overall accuracy and F1-Score, Logistic Regression excels at distinguishing ACVPs. Ensemble methods show promise but might require further tuning, especially for imbalanced datasets.

## IV. Conclusion

Peptide-based therapies hold immense potential beyond just antiviral drugs. They offer a wide range of therapeutic applications, including anti-cancer treatments [15]–[17]. This research highlights the promise of antiviral peptides (AVPs), specifically anti-coronavirus peptides (ACVPs), as a new weapon against coronaviruses. Traditional methods for finding ACVPs are slow and expensive. To address this, we introduced AntiCPs-CompML, an automated comprehensive machine learning-based method for efficient and accurate ACVP prediction with the application of LazyPredict Library. AntiCPs-CompML achieved high accuracy, paving the way for faster ACVP discovery and development of novel coronavirus treatments. This advancement has the potential to significantly improve human health and our ability to combat future viral threats.

## Notes

This work was supported in part by the National Research Foundation of Korea (NRF) grant funded by the Korea government (MSIT) (No. 2020R1A2C2005612) and (No. 2022R1G1A1004613) and in part by the Korea Big Data Station (K-BDS) with computing resources including technical support.

http://public.aibiochem.net/peptides/FEOpti-ACVP/

### Competing Interest Statement

The authors have declared no competing interest.

